# Omicron Spike Protein Is Vulnerable to Reduction

**DOI:** 10.1101/2023.01.06.522977

**Authors:** Zhong Yao, Betty Geng, Edyta Marcon, Shuye Pu, Hua Tang, John Merluza, Alexander Bello, Jamie Snider, Ping Lu, Heidi Wood, Igor Stagljar

## Abstract

SARS-CoV-2 virus spike (S) protein is an envelope protein responsible for binding to the ACE2 receptor, driving subsequent entry into host cells. The existence of multiple disulfide bonds in the S protein makes it potentially susceptible to reductive cleavage. Using a tri-part split luciferase-based binding assay, we evaluated the impacts of chemical reduction on S proteins from different virus variants and found that those from the Omicron family are highly vulnerable to reduction. Through manipulation of different Omicron mutations, we found that alterations in the receptor binding module (RBM) are the major determinants of this vulnerability. Specifically we discovered that Omicron mutations facilitate the cleavage of C480-C488 and C379-C432 disulfides, which consequently impairs binding activity and protein stability. The vulnerability of Omicron S proteins suggests a mechanism that can be harnessed to treat specific SARS-CoV-2 strains.

## Introduction

Since the beginning of the COVID19 pandemic, the SARS-CoV-2 virus has continued to evolve at a rapid pace. Numerous variants have been identified, including several variants of concern (VOCs): Alpha, Beta, Gamma, Delta and Omicron. The Omicron variant (B.1.1.529), characterized by improved transmissibility(Baker *et al*. 2022), but usually mild symptoms and less severe outcomes(Lewnard *et al*. 2022), was first identified in Botswana and South Africa in November of 2021(Viana *et al*. 2022). In a short period of time, it has evolved into numerous subvariants, among which BA.2, BA.4, BA.5 and their further derivatives have become the dominant strains in the world at the time of the writing of this report. In these variants, mutations occur in different areas of the genome and consequently may have varied impacts on virus biology as well as associated clinical outcomes. The mutations in spike (S) protein draw special attention due to the key roles of S protein in virus entry into host cells. Specifically, they affect its affinity to the ACE2(Cai *et al*. 2021) receptor, subsequently change virus transmissibility(Cai *et al*. 2021), and make a major contribution to immune evasion(Alter *et al*. 2021; Chen *et al*. 2021; Gobeil *et al*. 2021; Reynolds *et al*. 2022). Strikingly, an Omicron family S protein harbors more than 30 mutations(Viana *et al*. 2022), which may account for the fact that Omicron variants spread more easily than previous variants. More importantly, these mutations create resistance to antibodies raised by vaccination as well as most therapeutic antibodies(Cao *et al*. 2022; Carreño *et al*. 2022; Cele *et al*. 2022; Dejnirattisai *et al*. 2022; Iketani *et al*. 2022; Liu *et al*. 2022; Planas *et al*. 2022).

The S protein is an envelope protein which forms a homotrimer with its receptor binding domain (RBD) located at the top of the complex. The receptor binding motif (RBM), located at the tip of RBD, composes the surface directly contacting the ACE2 molecule(Lan *et al*. 2020; Shang *et al*. 2020). In the ectodomain of S protein, there are 30 cysteine residues. Structure studies(Cai *et al*. 2020; Walls *et al*. 2020; Wrapp *et al*. 2020) and mass spectrometry(Zhang *et al*. 2022) demonstrated that these cysteine residues form specific paired disulfide bonds (Expanded View Fig 1A). Interestingly, at least as known from the literature, no mutation in these cysteine residues has been observed in any variants (Fig EV1B), strongly supporting essentiality in virus biology. When considered alongside the capability of disulfide bonds to be reversibly reduced to free thiols, it seems probable that the S protein is potentially subject to redox regulation. Indeed, previous research suggested that changes in thiol/disulfide balance may affect the virus/host cell interaction in some viruses such as HIV(Fenouillet *et al*. 2007). Despite the fact that SARS-CoV S protein is relatively insensitive to redox change(Lavillette *et al*. 2006), recent studies showed that treatment of SARS-CoV-2 S protein with reducing compounds attenuated its activity toward ACE2-binding(Manček-Keber *et al*. 2021; Alonzi *et al*. 2022; Grishin *et al*. 2022; Khanna *et al*. 2022; Murae *et al*. 2022). However, most of these studies were based on the original Wuhan SARS-CoV-2 wild type (WT) strain or early variants. Whether the recent evolution of S protein affects its redox characteristics needs to be monitored, to help us fully understand the biology of SARS-CoV-2 and develop new methods of treatment and prevention.

In this study, we used a tri-part Nanoluciferase (tNLuc) based method called Neu-SATiN^30^ to measure the S/ACE2 interaction. We confirmed that S protein is susceptible to chemical reduction and found that different variants have varied sensitivities to reduction. Interestingly, Omicron variants BA.1, BA.2 and BA.4/5 exhibited remarkable vulnerability. We further found that the mutations in RBM are the major determinant to vulnerability of Omicron S proteins through facilitating cleavage of C480-C488 and C379-C425 disulfides. The cleavage leads to impairment in ACE2 binding and protein stability. Our observations suggest that the vulnerability of Omicron S protein can be harnessed for new approaches of COVID19 treatment and prevention.

## Results

### Different SARS-CoV-2 S protein variants present varied vulnerability to reduction

To evaluate the impact of chemical reduction on S protein function, we measured S/ACE2 binding after treating S protein with reducing compounds. Traditionally S/ACE2 binding affinity is measured with biophysical methods such as surface plasmon resonance (SPR), however these types of measurement are typically costly and require highly specialized equipment. We therefore used a simpler and more cost-effective approach, adapted from two tNLuc-based assays, <u>S</u>erological <u>A</u>ssay based on <u>T</u>r<u>i</u>-part <u>N</u>anoluciferase (SATiN)(Yao *et al*. 2021) and <u>Neu</u>tralization SATiN (Neu-SATiN)(Kim *et al*. 2022), previously developed by our group for measurement of antibodies against SARS-CoV-2 in human samples. In this approach, artificially trimerized S ectodomain protein or RBD protein is genetically tagged with the β10 tag of tNLuc. Similarly, ACE2 protein is attached to the β9 tag. S/ACE2 binding brings β9 and β10 into proximity and can be detected by addition of the third tNLuc component Δ11S, which results in reconstitution of a functional luciferase that produces measurable luminescence signal in the presence of the NLuc substrate furimazine (Fig 1A).

**Figure 1.**
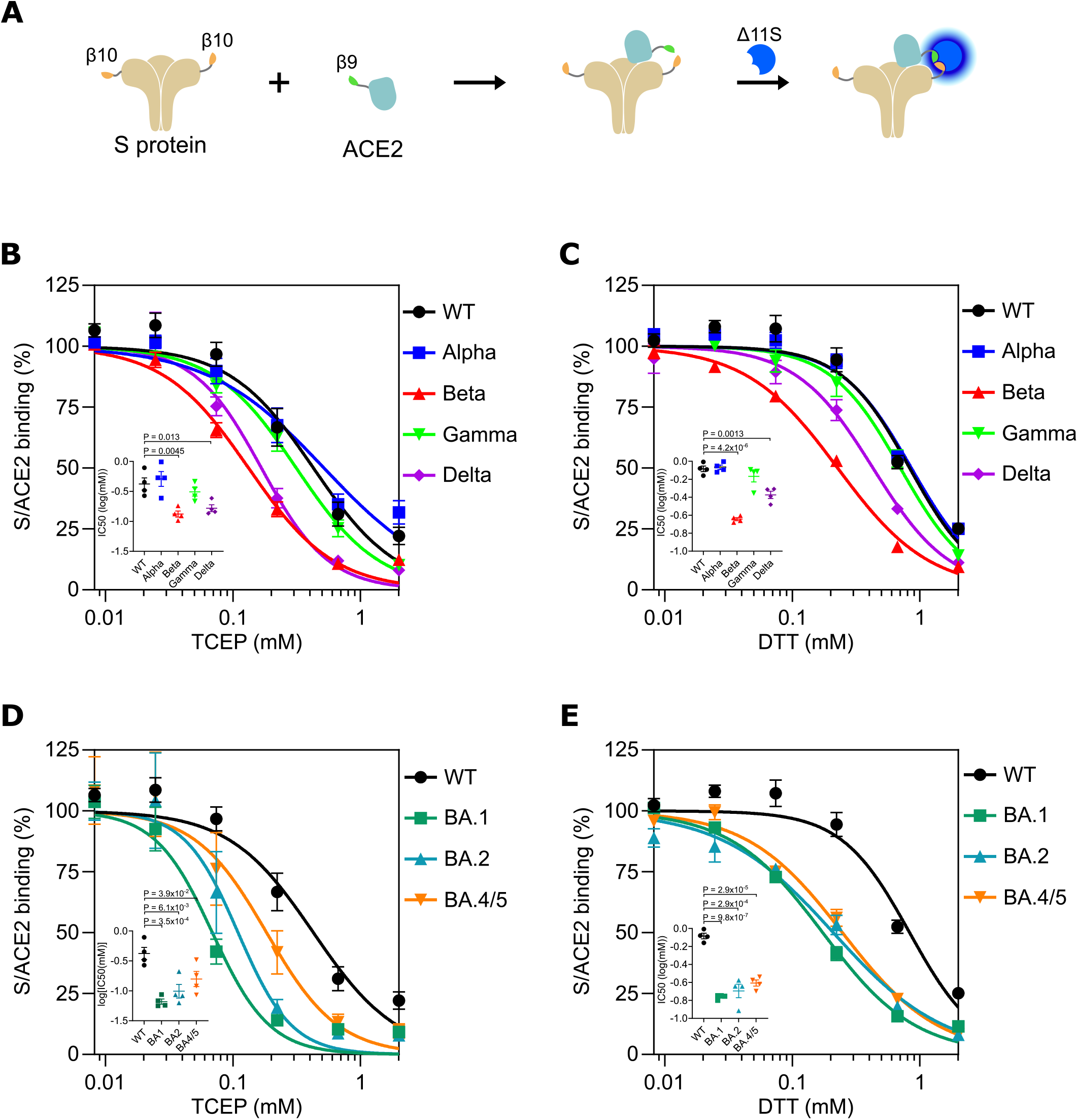
Different SARS-CoV-2 variant S proteins demonstrate varied vulnerabilities to chemical reduction. A Schematic diagram of the tNLuc-based S/ACE2 binding assay. B-E S proteins derived from early VOCs (Alpha, Beta, Gamma and Delta) (B,C) or Omicron variants (BA.1, BA.2 and BA4/5) (D,E) were treated with TCEP (B,D) or DTT (C,E) in various concentrations as indicated, followed by tNLuc-based S/ACE2 binding assay. The readings were normalized using corresponding S proteins with mock treatment. Data information: Results are presented as mean ± SEM of four independent experiments with two technical replicates in each experiment. Data were fitted into the inhibition kinetics model with normalized response and variable slope. Obtained kinetics are presented in average curves. IC50 values derived from each experiment are shown inset. P-values were calculated using a two-tailed t-test.

We first investigated if two commonly used strong reducing agents, tris-(2-carboxyethyl)phosphine (TCEP) and dithiothreitol (DTT), affect the binding ability of different S variants. Specifically, S proteins derived from parental strain (WT), Alpha, Beta, Gamma, Delta and Omicron (BA.1, BA.2 and BA.4/5) SARS-CoV-2 were incubated with varied amounts of TCEP or DTT (8 μM to 2 mM) followed by incubation with β9-ACE2 and luminescence measurement. BA.4 and BA.5 are interchangeably designated as BA.4/5 since S proteins in these two variants bear the same mutations. The results show that binding affinity to ACE2 of WT S protein was inhibited by both TCEP (Fig 1B) and DTT (Fig 1C) at a sub millimolar level, verifying previous observations that S protein can be inhibited by reduction. Among the earlier VOCs, Alpha and Gamma showed similar sensitivity to WT S protein. In contrast, Beta and Delta S proteins exhibited significantly enhanced sensitivity. For example, the Beta IC50 values of TCEP and DTT treatments are respectively 0.5 and 0.56 log unit lower than the corresponding values of WT S protein. Strikingly, all Omicron S proteins acquired even more marked sensitivity to reduction (Fig 1D, E). BA.1 exhibited the most significant shift of IC50, with 0.8 for TCEP and 0.68 for DTT log unit lower than WT S protein, followed by BA.2 and then by BA.4/5. Thus, we conclude that sensitivity to chemical reduction can be affected by different mutations in variant S proteins. Given that strong effects were observed on various Omicron variants and they are currently the dominant strains in the world, we decided to focus our further investigation on the BA.1, BA.2 and BA.4/5 S proteins.

TCEP and DTT are strong reducing agents, usually not suitable for administration to the human body for medicinal purposes. We thereby tested a panel of relatively mild antioxidants which have been approved by the FDA for usage in treatment of various diseases: reduced glutathione (GSH), N-acetylcysteine (NAC), N-acetylcysteine amide (NACA), bucillamine, cysteamine and WR1065 (Fig EV2). We observed that GSH and NAC did not exhibit any effects on WT S protein when administered at concentrations below 10 mM (Fig EV2A,B). In contrast, they inhibited BA.1, BA.2 and BA.4/5 S proteins with IC50 below 10 mM (Fig EV2A,B). NACA, bucillamine, cysteamine and WR1065 did have effects on WT S protein with IC50s in a range of 1-10 mM (Fig EV2C-F). Among them, bucillamine showed the most potent effect (IC50 ≈ 1 mM) (Fig EV2D). Consistent with the observations for TCEP and DTT, effects were significantly enhanced in Omicron variants, usually in the order of BA.1 > BA.2 > BA.4/5. In the case of NACA (Fig EV2C) which showed the best distinguishing between WT and Omicron variants, IC50 values of BA.1, BA.2 and BA.4/5 are respectively 0.42, 0.52 and 0.33 log units lower than WT. More strikingly, bucillamine showed 0.82, 0.70 and 0.40 log unit differences for BA.1, BA.2 and BA.4/5, respectively, from WT S protein, with IC50s of 149 μM, 198 μM and 402 μM, respectively, suggesting that bucillamine is a potent inhibitor (Fig EV2D). Thus, all these results support that Omicron variant S proteins are more vulnerable to chemical reduction.

### Omicron mutations in RBM are major determinants for the vulnerability to reduction

Omicron variants contain more than 30 mutations in their S proteins (Expanded View Fig 1A) compared to the original wild type (Wuhan) strain. We thereby sought to identify which mutations play causal roles in the vulnerability to reduction. Because they are distributed in different regions of the S protein, we first investigated which Omicron domain produces the most significant sensitivity using WT-Omicron subvariant (BA.1) S protein chimeras. An ‘S1-Omi’ construct, composed of an Omicron S1 fragment and a WT S2 fragment, displayed TCEP inhibitory kinetics massively shifted from that of WT S protein, reaching a position close to the Omicron curve (Fig 2A). This shift suggests that the alterations in S1 provide the major contribution to vulnerability. However, although the contributions by S2 mutations might be minor, they cannot be fully excluded since the S1-Omi curve did not reach inhibition fully equal to that of the Omicron S protein. In contrast, the inhibition kinetics of ‘NTD-Omi’, a WT S protein with replacement of the NTD Domain for that of Omicron, showed a similar profile to WT S protein, indicating that the alterations in NTD do not appear to be involved in sensitivity. The behavior of these two chimera constructs provided proof that changes in regions between NTD and S2 play important roles, leading us to focus on the RBD. Failing to produce a recombinant S protein with Omicron mutations only in the RBD, we instead created the ‘RBM-Omi’ construct with Omicron RBM in the WT backbone. Strikingly, its inhibition profile resembled that of the ‘S1-Omi’ construct (Fig 2A), suggesting that the Omicron mutations in RBM are the major determinants to the reduction vulnerability of Omicron S protein.

**Figure 2.**
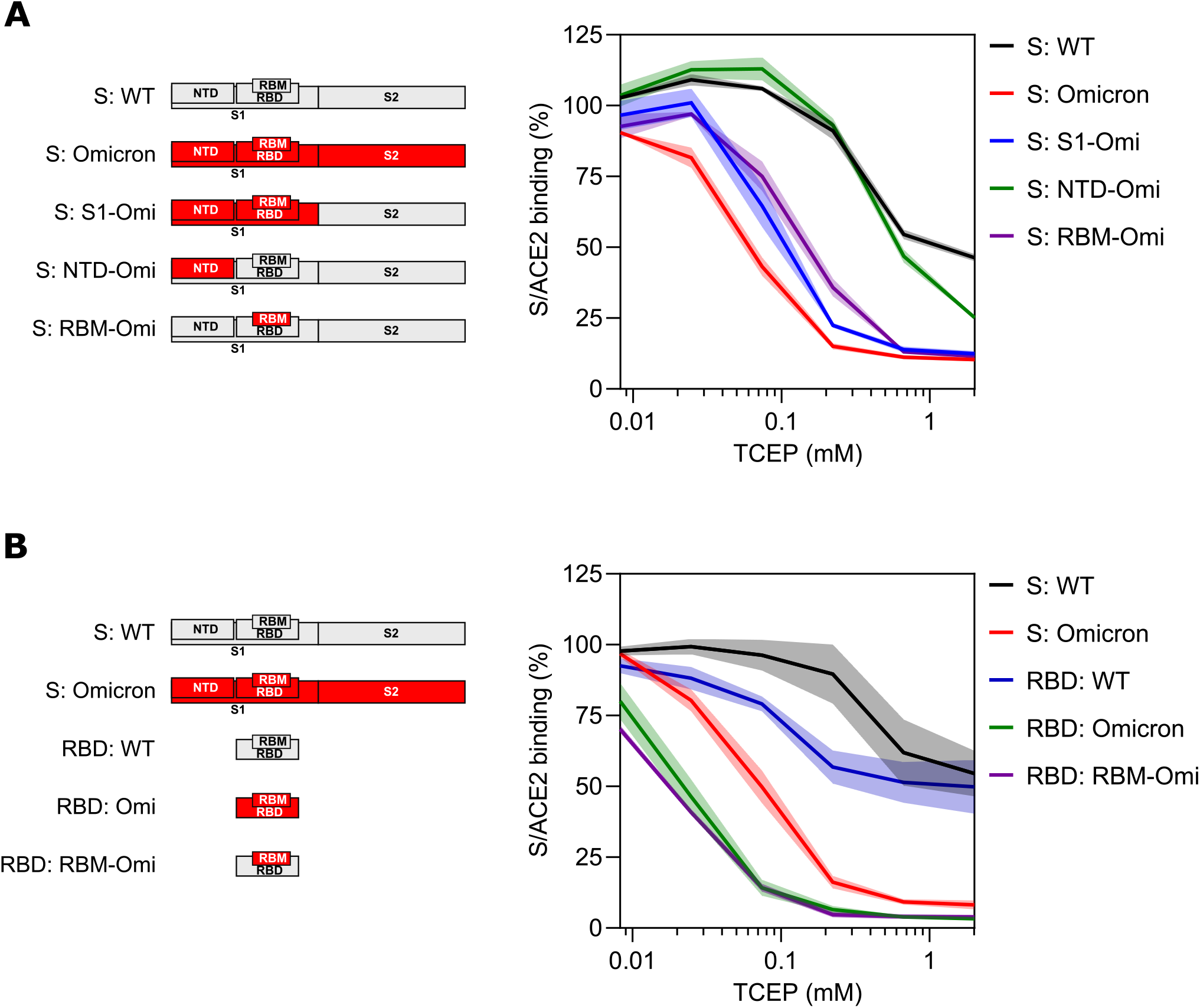
Omicron mutations in RBM are the major determinant in vulnerability to chemical reduction. A The impacts of omicron mutations were examined in WT/Omicron S protein chimeras. WT (S: WT), Omicron (S: Omicron), chimera of Omicron S1 and WT S2 (S: S1-Omi), chimera with Omicron NTD and WT backbone (S: NTD-Omi) or chimera with Omicron RBM and WT backbone (S: RBM-Omi) S proteins were pretreated with the indicated concentrations of TCEP followed by ACE2 binding assay. B Binding effects were further examined in RBD proteins. WT S protein (S: WT), Omicron S protein (S: Omicron), WT RBD (RBD: WT), Omicron RBD (RBD: Omi) or chimeric RBD with Omicron RBM on WT backbone (RBD: RBM-Omi) were pretreated with the indicated concentrations of TCEP followed by ACE2 binding assay. Data information: Readings were normalized with corresponding protein contructs with mock treatment. Results are presented as mean ± SEM of four independent experiments with two technical replicates in each experiment. SEMs are represented as error envelopes.

We further corroborated our findings through investigating the inhibition kinetics of TCEP with respect to RBD/ACE2 binding. Using RBD instead of the whole ectodomain of S protein allowed us to exclude any effects caused by other regions. Similar to the investigation of full S protein, RBD was pretreated with TCEP followed by the measurement of binding to ACE2. The TCEP inhibition profile of WT RBD showed a slight shift from that of WT S protein but maintained a similar baseline at high TCEP concentrations (Fig 2B), demonstrating its relative resistance to reduction. In contrast, the inhibition profile of RBD protein with only Omicron (BA.1) mutations in the RBM (‘RBM-Omi’ construct) displayed a massive shift from that of WT RBD, even further from Omicron S protein (Fig 2B), confirming the major role of the RBM in Omicron induced vulnerability to reduction. Interestingly, full Omicron RBD displayed an almost identical inhibition profile as the ‘RBM-Omi’ construct (Fig 2B), indicating that other mutations in RBD are at most limitedly involved in the regulation. We also asked if the involvement of Omicron RBM mutations is common in different Omicron variants. This question was answered by testing the RBM mutants in RBD proteins derived from BA.1 (same as RBM-Omi in Fig 2), BA.2 and BA.4/5 (Fig EV3). All of them demonstrated enhanced sensitivities to TCEP compared to WT protein but to different extents, consistent with the results obtained using their full-length S protein forms (Fig 1D).

Next, we sought to further narrow down the involved mutations within the RBM. The Omicron (BA.1) RBM harbors 10 substitution mutations (Fig 3A) compared to the S protein of the original wild type (Wuhan) strain, which we grouped into three clusters in the linear RBM sequence, cluster A (N440K, G446S), cluster B (S477N, T478K, E484A) and cluster C (Q493K, G496S, Q498R, N501Y, Y505H). Their roles were examined through both loss-of-function and gain-of-function construct design approaches (Fig EV4). We first prepared several loss-of-function mutants on either Omicron S protein backbone or RBM-Omi construct (RBD protein with Omicron RBM), in which an individual RBM cluster was reverted to its WT sequence (‘reverse mutant’). S protein with cluster C reverted was not included due to a failure in producing the protein. Compared to the WT, the cluster B reverse mutants (Omi-B) acquired a partial recovery of resistance to reduction in both full S protein and RBD protein contexts (Fig 3B,C). Reverse mutants of cluster A (Omi-A) or C (Omi-C) had no significant effect in RBD protein constructs (Fig 3C); Omi-A produced a moderate change in full S protein construct (Fig 3B). These observations suggest that cluster B plays a more important role than clusters A and C but is not sufficient to bring full vulnerability to reduction. This conclusion was further confirmed by gain-of-function mutants, specifically WT S proteins or RBD proteins harboring Omicron mutations in an individual RBM cluster. Notably, Omicron mutations in cluster C (C-Omi) either had no effect in S protein (Fig 3D) or produced the least sensitivity to TCEP in RBD protein (Fig 3E). Mutations in cluster A (A-Omi) and B (B-Omi) produced similar significant sensitivity to TCEP but did not reach to the levels of Omicron S protein or Omicron RBD protein. Putting these results together, we conclude that the importance of these clusters in determining the vulnerability to reduction can be listed in an order as B>A>C, however, all of them are required for full sensitivity.

**Figure 3.**
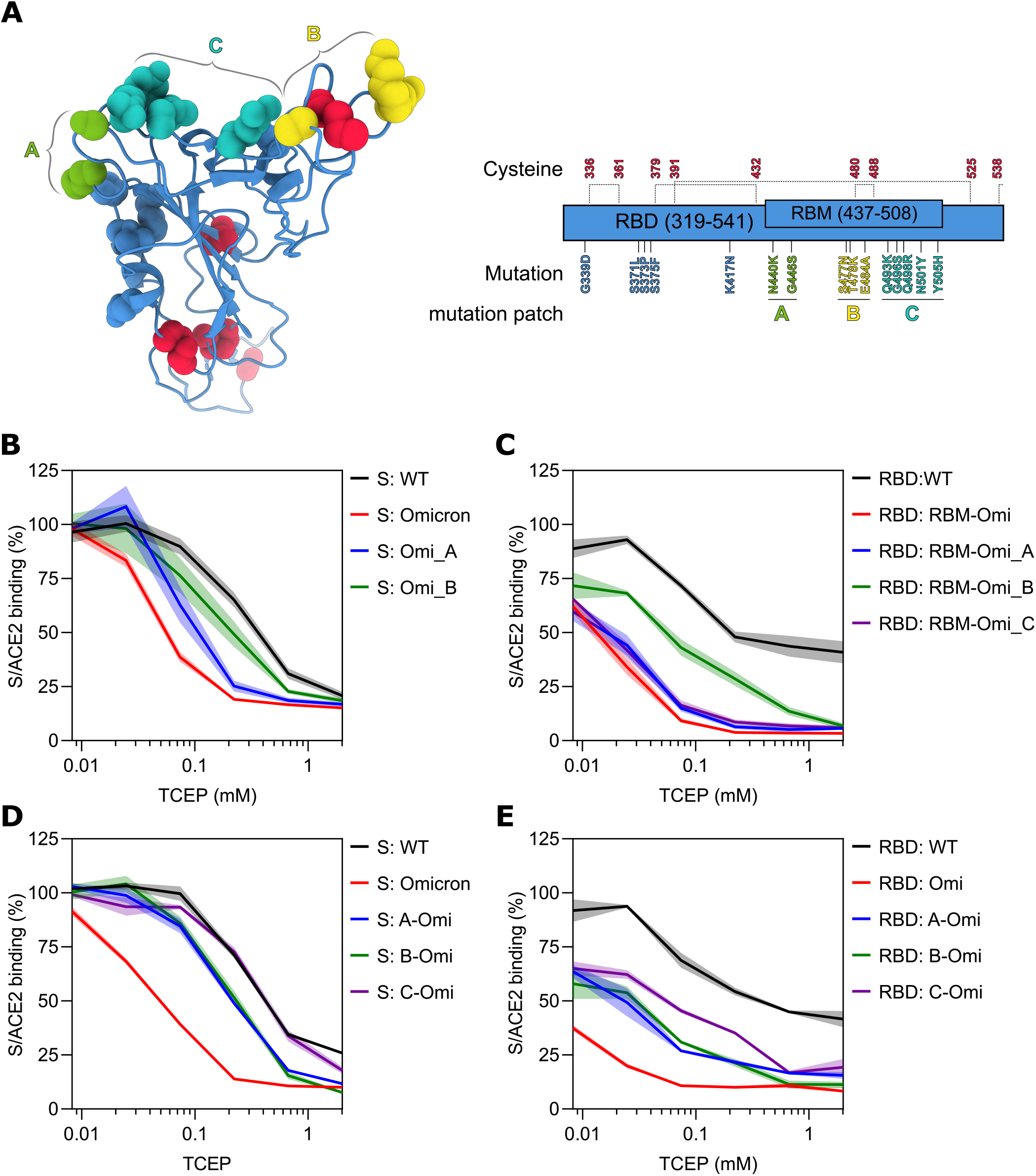
Detailed analysis of the importance of Omicron RBM mutations in the vulnerability to chemical reduction. A 3D and linear structure of the Omicron RBD. Cystines are highlighted in red and represented as red spheres. Mutations are also represented as spheres and shown in different colors based on their grouping into three clusters. B-C Analysis of loss-of-function mutants. In B, WT and Omicron S proteins and S proteins of the Omicron backbone with cluster A or B mutations reverted to WT sequence (S: Omi_A or S: Omi_B) were analysed. In c, WT RBD protein, WT RBD carrying Omicron RBM (RBD: RBM-Omi), and WT RBD protein carrying Omicron RBM with clusters A, B or C reverted back to WT sequence (RBD: RBM-Omi_A, RBD: RBM-Omi_B or RBD: RBM-Omi_C) were analysed. Protein constructs are listed in Fig EV4. D-E Analysis of gain-of-function mutants. In D, WT and Omicron S proteins, and S proteins of WT backbone with an RBM cluster mutated to its Omicron sequence (S: A-Omi, S: B-Omi or S: C-Omi) were analyzed. In E, WT RBD protein and Omicron RBD protein (RBD: Omi), alongside RDB proteins with WT backbone and an RBM cluster mutated to its Omicron sequence (RBD: A-Omi, RBD: B-Omi or RBD: C-Omi) were analyzed. Proteins in B-E were pretreated with the indicated concentrations of TCEP followed by ACE2 binding assay. Protein constructs are listed in Fig EV4. Data information: Readings were normalized with corresponding S or RBD proteins with mock treatment. Results are presented as mean ± SEM of four independent experiments with two technical replicates in each experiment. SEMs are represented as error envelopes.

### C480/C488 and C379/C432 disulfides in Omicron S protein are susceptible to reduction

To determine how Omicron mutations cause vulnerability, we sought to identify which disulfide bonds in the RBD are subject to chemical reduction-mediated cleavage. Because we have concluded that mutations in the RBM play major roles, we focused the investigation on the RBD using RBD protein constructs.

There are nine cysteine residues in the RBD, forming 5 disulfide bonds (Fig 3A). Among them, the C480-C488 disulfide is at the tip of RBD, flanked by cluster B Omicron mutations, while disulfide C379-C432 is in the middle of RBD. In contrast, disulfides C336-C361 and C391-C525 are at the bottom of the RBD structure. Also located at the bottom is C538, forming a disulfide bond with C590 outside of the RBD and therefore not included in the RBD construct. Thus, the redox state of C538 was left unexplored in this study.

We used an approach of chemical labelling coupled with mass spectrometry (MS) to identify the reduced cysteines (Fig 4A). WT and Omicron RBD proteins were pretreated with TCEP at a concentration (0.1 mM) that inhibits >80% activity of Omicron RBD but only has marginal effect on WT RBD. Control samples were mock treated. This was followed by alkylation with iodoacetamide (IAM), which adds a carbamidomethyl group to the free thiol group of reduced cysteine. After trypsin digestion, the identities and quantities of cysteine-modified peptides were resolved by MS (Fig 4A) with the reduced cysteines detected by alkylation. In total, nine distinct alkylated cysteine containing peptides were identified in all samples, covering eight different cysteines (Fig 4B). Since some cysteine residues (C361, C432 and C480/C488) are surrounded with mutated amino acids in Omicron peptides, each of them appears in two different peptides in WT and Omicron samples. Excluding C538, seven out of eight total target cysteine residues were analyzed in this experiment. In the WT RBD samples (Fig 4B), the alkylation signals were usually low or with some even below the detection limit: no reduction of C379, C480 and C488 was observed at basal state, fully consistent with previous work(Shi *et al*. 2022).Upon TCEP treatment, most of the cysteines did not show increase in reduction, indicating that disulfides in WT RBD are usually relatively inert in response to chemical reduction. In contrast, alkylation signals of many cysteine residues in the Omicron sample maintained a relatively high basal level (Fig 4B). This might suggest that disulfides in the Omicron RBD are less stable, consistent with the general thermodynamically less stable state of the Omicron RBD structure found in a previous study(Yin *et al*. 2022). Interestingly, TCEP treatment led to reduction on at least six Omicron cysteine residues. Among them, C432, C379 and C480/C488 showed respectively 8.3, 2.6 and 2.3 fold change relative to untreated cysteines, providing proofs that C379-C432 and C480-C488 disulfides are subject to cleavage. It should be noted that C432 or C480-C488 peptides are distinct in WT and Omicron samples and should not be directly compared. Nonetheless, the fact that they could not be detected in TCEP treated WT samples still suggests that they are relatively inert to reduction in WT RBD, confirming a previous observation by Shi et al(Shi *et al*. 2022). Therefore, C379-C432 and C480-C488 disulfides are highly susceptible to reduction in the Omicron RBD.

**Figure 4.**
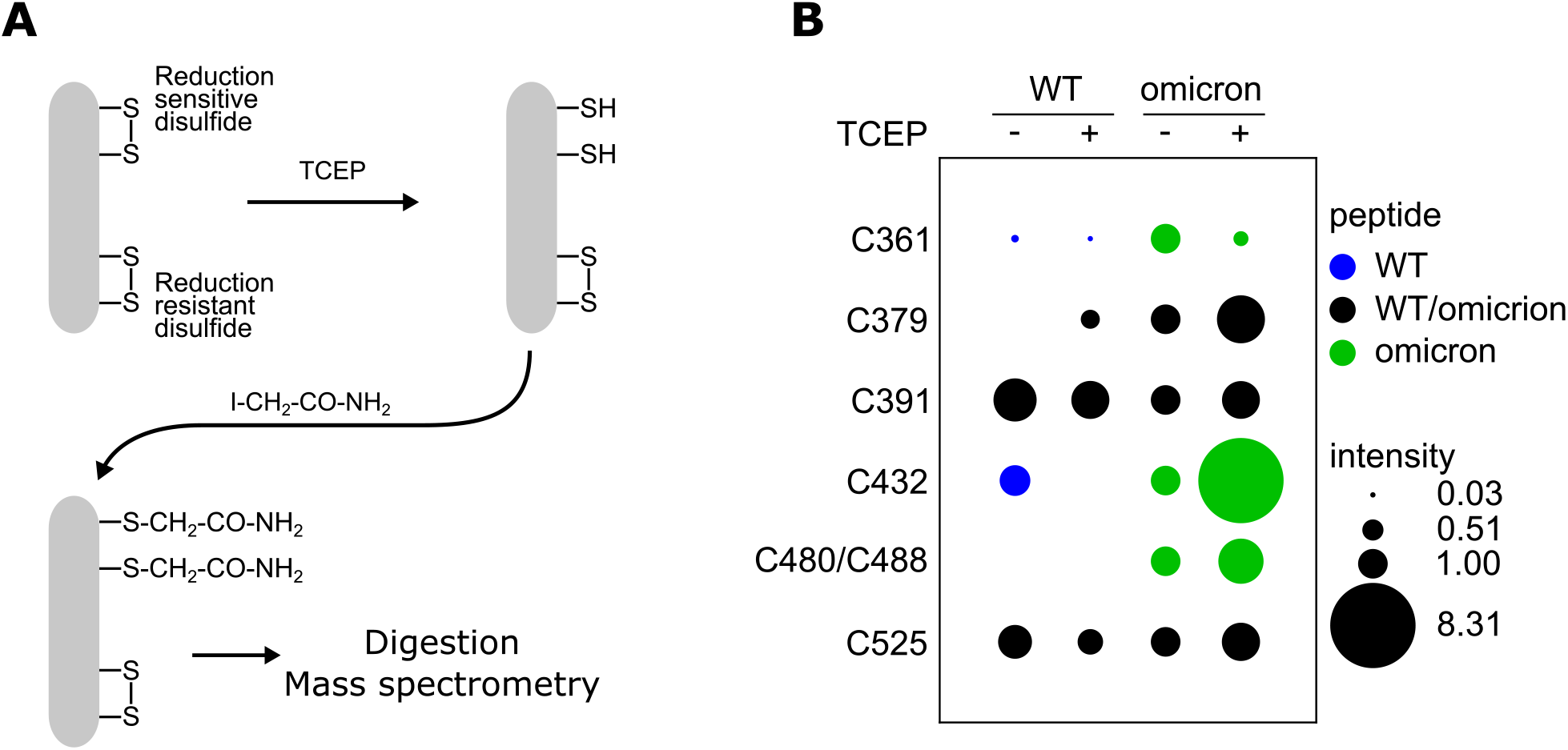
Mass spectrometry analysis of reduced cysteines in RBD. A Strategy of the MS analysis. B WT or Omicron RBD proteins were treated with 0.1 mM TCEP or mock treated and analyzed as described in A. The intensities of peptides containing alkylated cysteines derived from RBD proteins are presented as a dot plot. The data are normalized to the intensities of nontreated Omicron RBD sample. Black dots stand for common peptides appearing in both WT and Omicron RBD; blue dots stand for WT-specific peptides; green dots stand for Omicron specific peptides.

### The impacts of disulfide cleavage in Omicron RBD

To test how disulfide cleavage affects functionality of S protein, we created Omicron RBD proteins with C336S/C361S, C379S/C432S, C391S/C525S or C480S/C488S mutations to mimic breakage of disulfide bonds. We first measured the stability of these mutants. Expression constructs of these mutants and a plasmid expressing secreted *Cypridina noctiluca* luciferase(Wu *et al*. 2007) were transfected into HEK 293 cells. After 20 hours, the quantity of RBD protein in culture media was measured using the HiBiT system(Dixon *et al*. 2016) targeting the β10/HiBiT tag attached to the RBD protein, normalized by *Cypridina* luciferase reading (Fig 5A). Indeed, all CS mutants displayed loss of expression to varied extent, suggesting all the corresponding disulfide bonds are involved in stabilizing the protein structure. Disulfide bonds C379-C432, C391-C525 and C480-C488 may play more important roles given that the corresponding mutants showed about a 90% loss compared to Omicron RBD protein levels.

**Figure 5.**
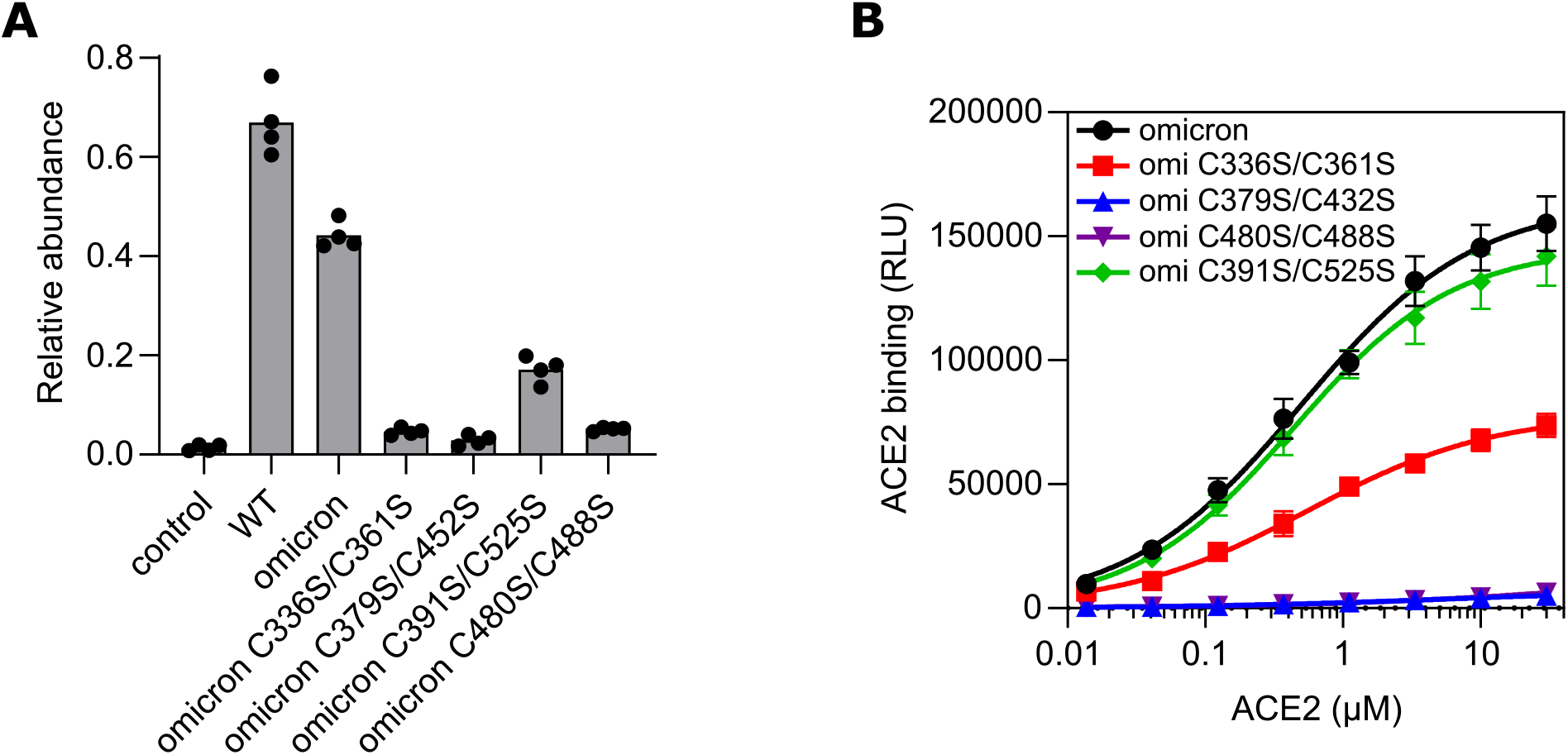
Impacts of disulfide cleavage in Omicron RBD. A Disulfide cleavage was mimicked by paired cysteine-to-serine mutation. Protein stability of the mutants was measured as the abundance of each protein in culture medium of the cells transiently expressing the protein. The results are normalized using co-expressed *Cypridina* luciferase. B The binding activity of each cysteine-to-serine mutant was measured using a tNLuc-based assay with different concentrations of ACE2. The results are presented as mean ± SEM of three independent experiments with two technical replicates in each experiment.

The impacts of disulfide cleavage on RBD binding to ACE2 were also investigated using the cysteine substitution mutants by profiling their binding kinetics with varied concentrations of ACE2 under the condition of equal amounts of RBD mutant proteins were used (Fig 5B). C391S/C525S mutant presented a profile almost matching Omicron RBD, indicating that the conformational change caused by cleavage of C391-C525 disulfide does not affect ACE2 binding. In contrast, ACE2 binding activity were both eliminated in C379S/C432S and C480S/C488S mutants, suggesting the key roles of C379-C432 and C480/C488 disulfides in maintaining Omicron RBD in a structure competent for ACE2 binding. Consistently, the importance of C480-C488 disulfide was also observed in WT RBD in a previous study using a C488A mutant(Murae *et al*. 2022). Interestingly, although C336S/C361S mutation caused a significant defect in RBD stability comparable to C379S/C432S and C480S/C488S mutants (Fig 5A), it only led to about 50% reduction in its binding activity (Fig 5B), suggesting that the correspondent conformational change contributes more to damage to protein stability than to loss of activity. Combining the MS results with functionality data together, we conclude that Omicron mutations make C379-C432 and C480-C488 disulfides susceptible to decreases in environmental redox potential. Their reduction induced cleavage then leads to decreases in both the activity and stability of Omicron S protein.

## Discussion

In this study, we found that Omicron S proteins are highly vulnerable to chemical reduction. To clarify the mechanism, we first defined the causal relationship of Omicron mutations and this vulnerability. Using a range of WT/Omicron chimera S proteins, we found that Omicron alteration in the RBM is the major determinant of the vulnerability. Specifically, the mutations surrounding C480-C488 disulfide provide a major contribution, although other RBM mutations are also required to produce full vulnerability. Next, we explored exactly which disulfides are susceptible to reduction in the Omicron variants. Using MS analysis, we first confirmed that disulfides C480-C488 and C379-C432 are relatively inert in WT RBD, consistent with results described in a previous report(Shi *et al*. 2022). The relative inertness was preserved in WT S protein even when treated with a low TCEP concentration. However, the same amount of TCEP induced markedly increased cleavage of C480-C488 and C379-C432 disulfides in Omicron variants, providing proof of their high sensitivity to reduction. The question of how exactly these Omicron mutations lead to vulnerability, however, still needs to be clarified. While our work in this report did not dive into a detailed mechanistic study in this regard, there are useful clues both in our results and in published data. For example, the local Omicron RBD structure seems to be thermodynamically less stable(Yin *et al*. 2022) although the overall structure of Omicron trimer S protein is in a more compact conformation(Cui *et al*. 2022; Gobeil *et al*. 2022). Thus, local mobility may make disulfides entropically less stable and more accessible to reducing agents. Additionally, the E484A and T478K mutations cause a negative to positive shift in the electrostatic micromilieu of the C480-C488 disulfide, a change which may also affect the redox potential(Jensen *et al*. 2009; Bechtel and Weerapana 2017). These hypotheses both merit further examination in the future to establish their potential validity.

A fundamental question relevant to the Omicron vulnerability is what the consequences are of disulfide cleavage. Like many viruses, cysteine residues in the ectodomain of SARS-CoV-2 S protein form disulfides (cystines), which are maintained by the relatively oxidizing extracellular milieu and infection-induced oxidative stress(Suhail *et al*. 2020). These disulfides are supposed to play key roles in functions such as proper folding, protein stability and binding activity toward ACE2, which are supported by molecular dynamics studies based on simulation of full disulfide cleavage in WT RBD(Fossum *et al*. 2022; Grishin *et al*. 2022). Considering simultaneous reduction of all four disulfides rarely occurs in the nature, it might be more relevant to characterize individual disulfides. However, different computational studies of single disulfide cleavage showed controversial results(Grishin *et al*. 2022; Shi *et al*. 2022). Thus, experimental approaches could provide more reliable proof. Here, we created mutants of paired cysteine-to-serine substitution to mimic reduction of each disulfide in the Omicron (BA.1) RBD protein. We conclude that C480-C488 and C379-C432 disulfides play key roles in S protein functionality since disruption of either disulfide severely impaired both protein stability and binding activity. As C480 and C488 are in the RBM, we reason that the disulfide may support a local surface geometry competent for ACE2 binding. The detailed role of C379-C432 at the atomic level needs further investigation although a previous computational study has provided some hints that its cleavage can cause massive structure change(Shi *et al*. 2022). In contrast to our results with C480-C488 and C379-C432, we found that disruption of C391-C525 had very limited effect, while disruption of C336-C361 severely destabilized the protein but only mildly attenuated its binding activity. Given the long-range distance between the RBM and these two disulfides, we rationalize that the effects of conformational changes caused by C336-C361 or C391-C525 cleavage may not be transferred to the RBM region even though they can bring about some global changes. It should be noted that previous investigations were performed in the context of full WT S proteins. Here we study the importance of C480-C488 and C379-C432 in Omicron S protein. Whether the intactness of these two disulfides is equally important in WT and Omicron S proteins merits more elaboration.

Previously the effects of reduction have been demonstrated in both cellular models(Manček-Keber *et al*. 2021; Alonzi *et al*. 2022; Grishin *et al*. 2022; Khanna *et al*. 2022; Murae *et al*. 2022) and mouse models(Khanna *et al*. 2022). Here, we verified the same phenomenon using an *in vitro* method and elaborated on the involved mechanism. All these studies suggested that this sensitivity can be potentially harnessed for fighting COVID-19. However, most of previous studies were based on WT S protein and the observed effects were limited, judging by the fact that high concentrations of reducing compounds were required. Thus, the true effects of corresponding clinical interference need to be verified. Notably, two reducing agents, bucillamine (https://clinicaltrials.gov/ct2/show/NCT04504734) and cysteamine (https://clinicaltrials.gov/ct2/show/NCT05212662), are under clinical trials and the results are awaiting release. With the dominance of Omicron variants, it becomes important to evaluate their sensitivity to chemical reduction. Our study clearly demonstrated a significant vulnerability of Omicron variant compared to WT, which was also mentioned by two previous studies(Alonzi *et al*. 2022; Khanna *et al*. 2022). Thus, more potent effects in the Omicron variant can potentially be achieved with the same concentration of reducing agent. Our mechanistic study further provided a rationale for reduction-based treatment for Omicron-induced disease. It should be noted that the lowest effective concentration of the reducing agents is above 100 μM, which is difficult to reach in the human body if delivered through oral or intravenous administration. Thus, direct delivery to respiratory tract through aerosol nebulization may be a better approach. In addition, whether the vulnerability is involved in the life cycle of Omicron virus in human body and in the clinical outcome of human infection also requires investigation in the future.

Our study demonstrated that the Omicron mutations around the C480-C488 disulfide, S477N, T478K and E484A, play important roles in causing the observed vulnerability. Based on the fact that these are signature mutations that are preserved in all Omicron subvariants, we rationalize that the vulnerability likely exists in all Omicron variants. Here we confirmed in BA.1, BA.2, BA.4 and BA.5 variants, however, considering the rapid rate of evolution of SARS-CoV-2, verification in more recent Omicron variants is merited.

Here, we reported a marked vulnerability to chemical reduction in Omicron S proteins. The involved mechanism may lay as a foundation for treatment and prevention of Omicron related infections.

## Materials and Methods

### Chemicals

Chemicals used in this study are listed as follow. TCEP: BioShop TCE101; DTT: BioShop DTT001; Reduced glutathione: BioShop GTH001; N-Acetyl-L-cysteine: Sigma-Aldrich A9165; N-acetylcysteine amide: Selleckchem 38520-57-9; Bucillamine: Toronto Research Chemicals B689375; Cysteamine: Sigma-Aldrich M9768; WR1065 Sigma-Aldrich W2020; Iodoacetamide: BioShop IOD500.

### Plasmids and recombinant proteins

Plasmids and recombinant proteins were prepared as previously described(Yao *et al*. 2021; Kim *et al*. 2022). Mutations were created using Gibson assembly (NEB E2611).

### Effects of reducing compounds on S/ACE2 binding

S or RBD protein was diluted in buffer A (Tris pH7.5 20 mM, Triton X-100 0.1%, EDTA 2 mM, NaCl 100 mM and fatty acid free BSA 0.05%) with final concentration of 5 nM and was incubated with reducing compounds at different concentrations at 37^0^C for 30 minutes for DTT or TCEP or 60 minutes for GSH, NAC, NACA, bucillaime, cysteaime and WR1065. Subsequently 4 μl of the treated sample was mixed with 36 μl of buffer B (Tris pH7.5 20 mM, Tween 20 0.1%, EDTA 2 mM, NaCl 15 mM, BSA (fatty acid free) 0.05%) supplemented with furimazine 25 μM, ACE2 0.5 μM and Δ11S 0.2 μM, followed by incubation at room temperature for 60 minutes. Luminescence was read using a microplate luminesce reader. IC50s were obtained by fitting the data into a normalized inhibition kinetics with variable slope using GraphPad 9.5.0.

### Identification of reduced cysteine by Mass spectrometry analysis

WT or Omicron RBD protein (5 μM in 20 μl) was mixed with 20 μl buffer A containing TCEP (0.2 mM) or blank and incubated at 37^0^C for 30 minutes. IAM solution (25 mM 10 μl) was then added and incubated in dark at room temperature for 30 minutes. Next, DTT (60 mM in 10 μl) solution was added and incubated in dark at room temperature for 60 minutes. Mass spectrometry analysis was performed as described. Briefly, the protein was further digested by trypsin over night. The digested samples were dried and re-suspended in 200μL 80% acetonitrile/2% formic. The peptides were injected into C18 nano-column on an EASY-nLC (Proxeon) coupled with an Orbitrap Velos mass spectrometer (Thermo Fisher Scientific). A ninety-minute gradient was used for separation at a flow rate of 300nL/minutes. Sixteen MS/MS collision induced dissociation (CID) data-dependent scans were acquired simultaneously with one full scan mass spectra with 60000 resolution and 1E6 AGC. MaxQuant (version 2.0.3.0) was used to search Uniprot/SwissProt human protein database with 20,600 human proteins modified to contain BSA and trypsin, and with variable modification of methionine oxidation, protein N-terminal acetylation and cysteine carbamidomethylation as variable modificatios and cysteine carbamidomethylation as a fixed modifications. Up to 1% false discovery rate was used for both peptides and proteins.

### Stability assay

Plasmids of *Cypridina noctiluca* luciferase and Omicron RBD or its mutants were co-transfected (9:1) into HEK293 cells using X-tremeGENE 9 (Millipore-Sigma XTG9-RO). After 16 hours, RBD proteins in culture media were measured in Nano-Glo buffer (Promega N1110) supplemented with Nanoluciferase substrate and 11S protein according to the manual. Cypridina luciferase vargulin was measured by mixing 5 μl medium with 20 μl phosphate buffer saline supplemented with vargulin 3 μM followed by luminescence reading. The amount of RBD proteins were normalized with Cypridina luciferase activity.

### RBD/ACE2 affinity assay

To measure the affinity of Omicron RBD cysteine-to-serine mutants to ACE2, varied concentrations of ACE2 was mixed with the mutant RBD protein (final concentration 0.8 nM) in buffer A. An aliquot of the mixture (10 μl) of the mixture was mixed with 30 μl of buffer B supplemented with furimazine 25 μM and Δ11S 0.2 μM. Luminescence was read after 2-hour incubation at room temperature.

## Acknowledgements

We thank the members of Igor Stagljar’s lab for their suggestions. This work was supported by the Toronto COVID-19 Action Fund (Connaught Award #0000313897) and Temerty Knowledge translation grant.

## Author Contributions

Z.Y. conceptualized the work. Z.Y., I.S. and P.L. designed the experiments. Z.Y., B.G., E.M., H.T., J.M., A.B. and H.W. performed experiments, S.P. analysed the MS data, Z.Y. analysed and interpretated the data. Z.Y., J.S. and I.S. wrote the manuscript.

## Legend

**Expanded View Figure 1.**
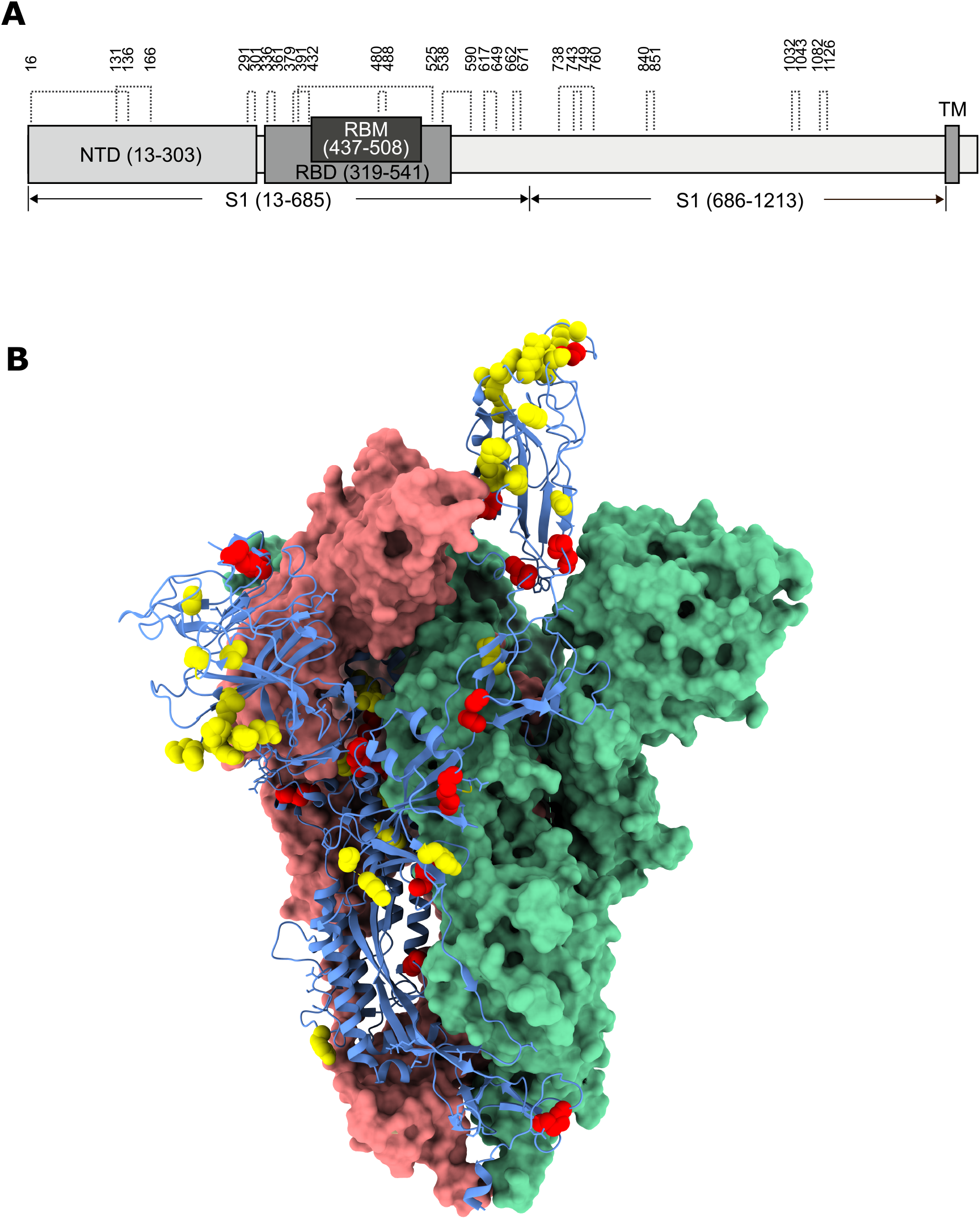
Disulfide bonds in SARS-CoV-2 spike protein. A Schematic representation of cysteine residues and disulfides located in different parts of the S protein. B Spatial location of disulfides in Omicron S protein. One protomer is presented as a blue cartoon model with cysteines highlighted as red spheres and mutation sites as yellow spheres. It should be noted that no mutation at cysteine residues has been identified in any variant, as exemplified here in Omicron S protein.

**Expanded View Figure 2.**
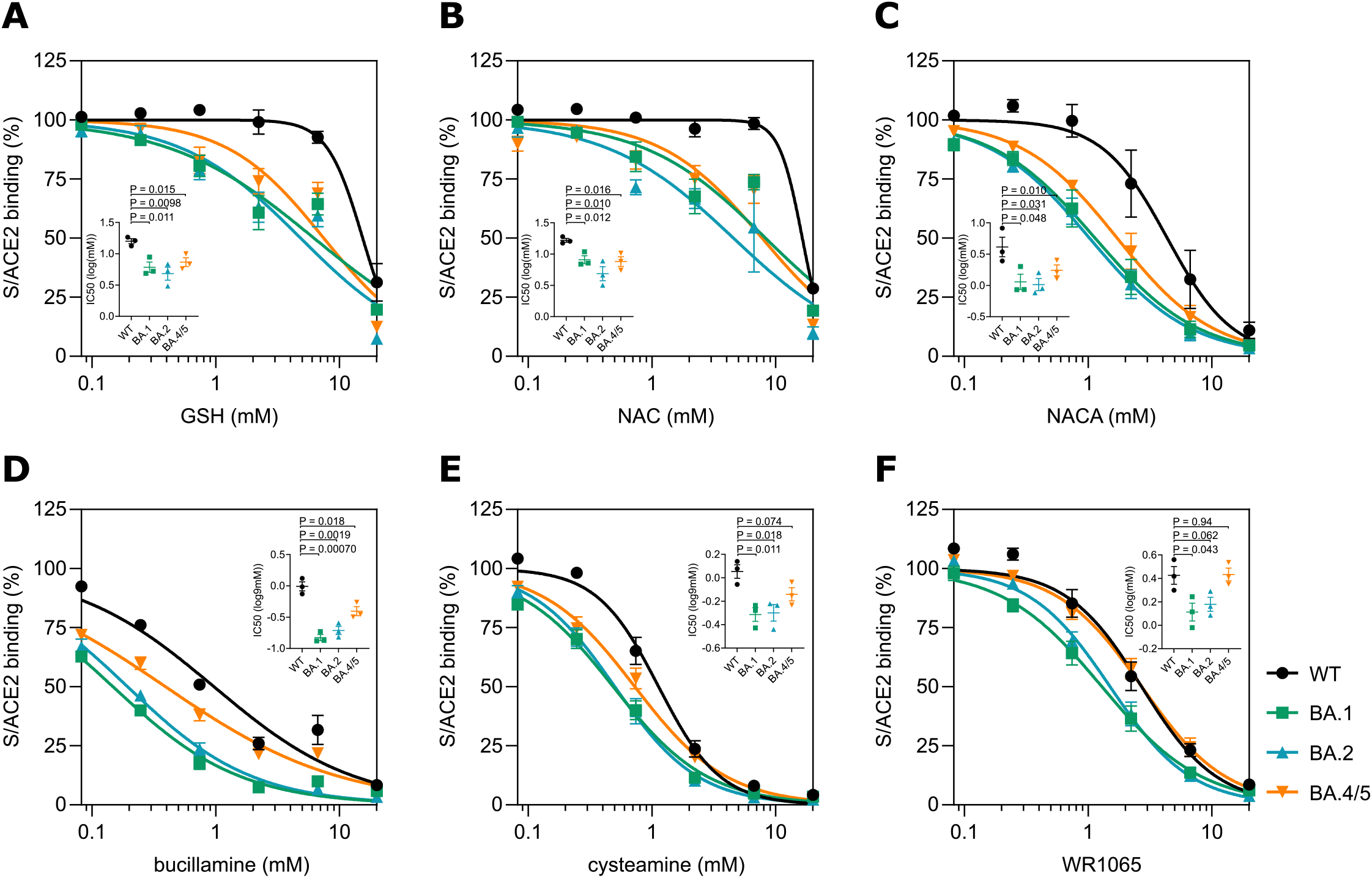
Effects of FDA-approved antioxidants on Omicron S proteins. S proteins derived from WT virus or Omicron variants (BA.1, BA.2 and BA4/5) were treated with different FDA-approved antioxidants in various concentrations as indicated, followed by tNLuo-based S/ACE2 binding assay. The readings were normalized using corresponding S proteins with mock treatment. Results of four independent experiments with two technical replicates in each experiment for each S protein are presented as mean ± SEM. Data were fitted into the inhibition kinetics model with normalized response and variable slope. Obtained kinetics were presented in average curves. IC5O values derived from each experiment are shown inset. P-values were calculated using a two-tailed t-test.

**Expanded View Figure 3.**
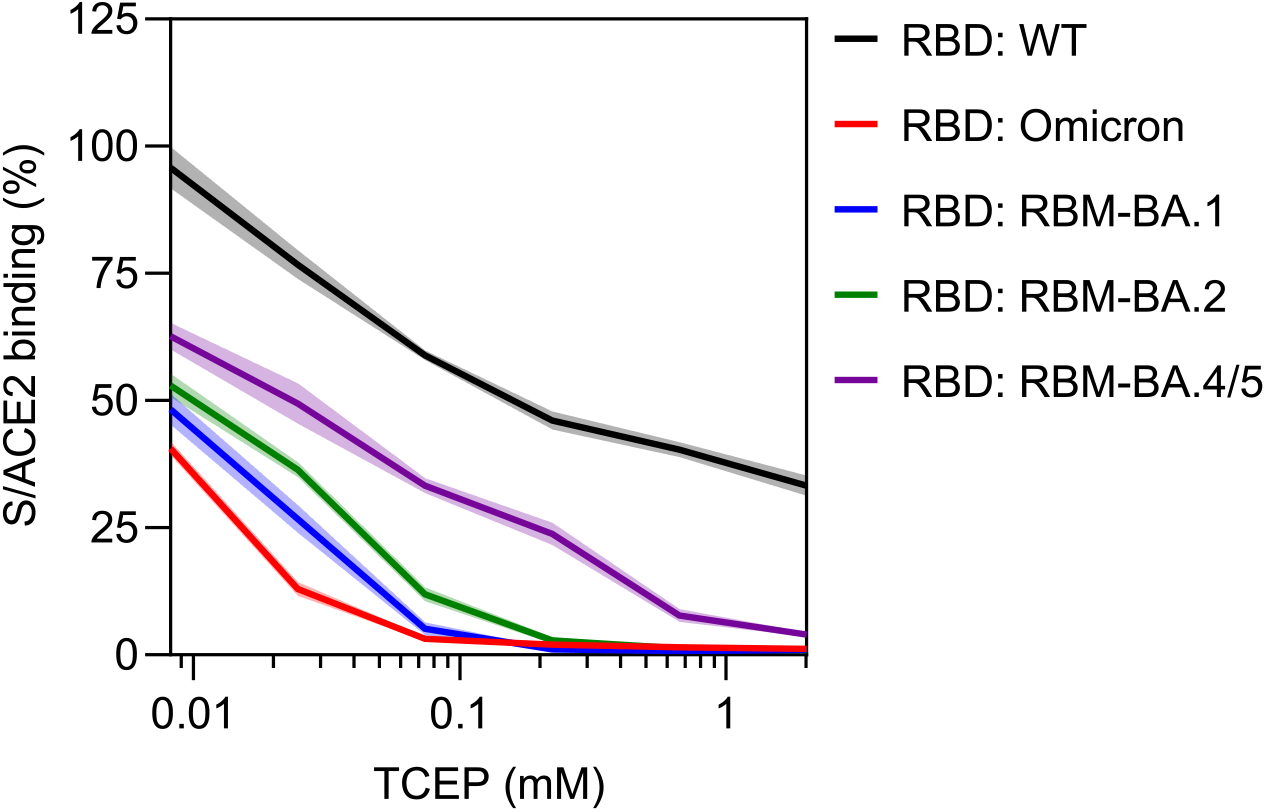
Verification of the important role of RBM mutations in vulnerability to reduction across different Omicron variants. Together with WT and Omicron/BA.1 RBD proteins (RBD: RBM-BA.1), RBD proteins with the RBM mutated to match different Omicron variants (RBD: RBM-BA.2, RBD: RBM-BA.4/5) were pretreated with the indicated concentrations of TCEP followed by ACE2 binding assay. Readings were normalized with corresponding RBD proteins with mock treatment. Results are presented as mean ± SEM of four independent experiments with two technical replicates in each experiment. SEMs are represented as error envelopes.

**Expanded View Figure 4.**
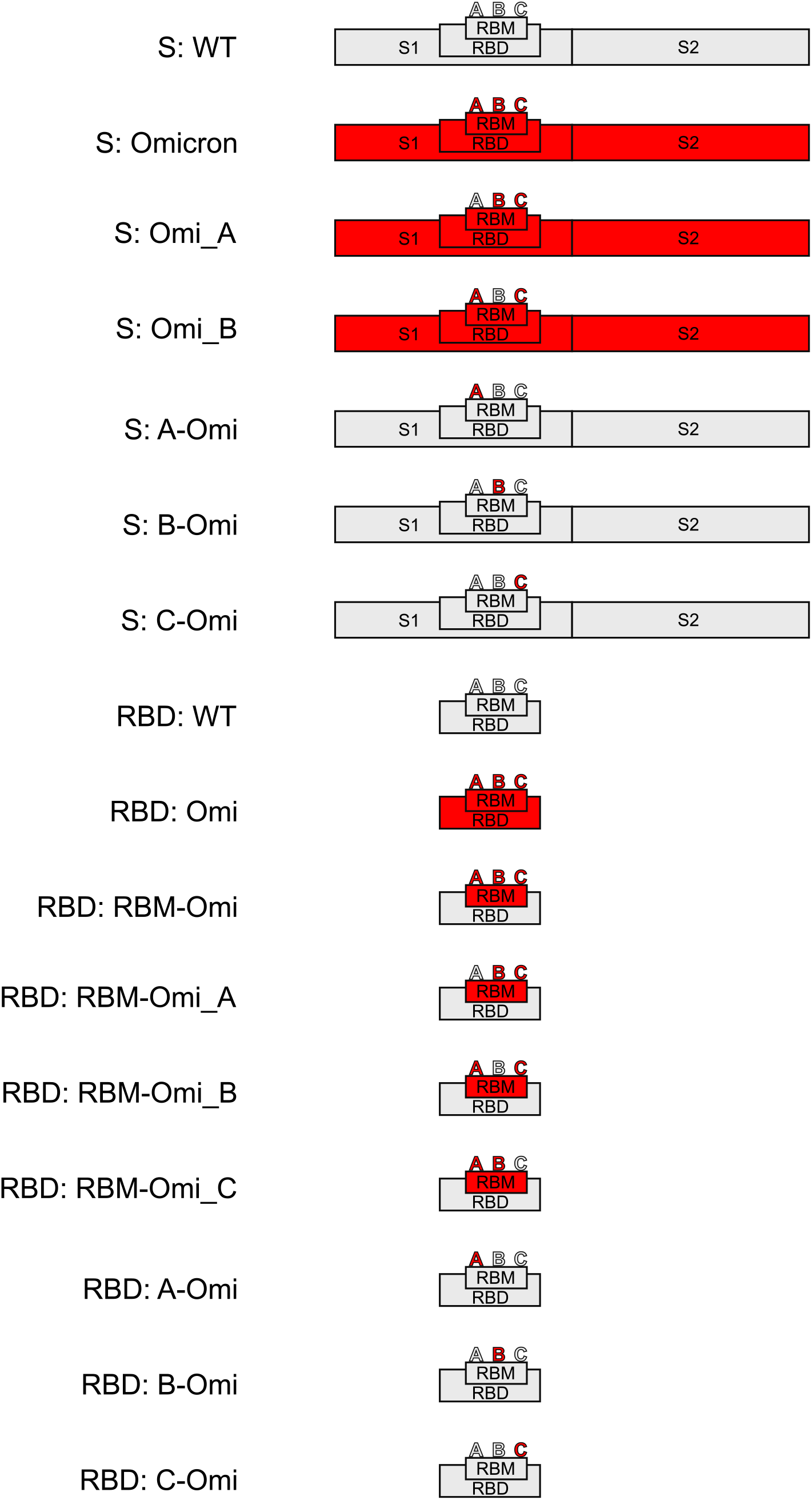
Schematic representation of S or RBD mutants used in Figure 3. WT sequences are highlighted in grey and Omicron sequences are in red.

## Reference

Alonzi, T., Aiello, A., Repele, F., Falasca, L., Francalancia, M., Garbuglia, A.R., Delogu, G., Nicastri, E., Piacentini, M., and Goletti, D. (2022) ‘Cysteamine exerts in vitro antiviral activity against the SARS-CoV-2 Delta and Omicron variants.’, Cell death discovery, 8(1), 288, available: https://doi.org/10.1038/s41420-022-01080-8.

Alter, G., Yu, J., Liu, J., Chandrashekar, A., Borducchi, E.N., Tostanoski, L.H., McMahan, K., Jacob-Dolan, C., Martinez, D.R., Chang, A., Anioke, T., Lifton, M., Nkolola, J., Stephenson, K.E., Atyeo, C., Shin, S., Fields, P., Kaplan, I., Robins, H., Amanat, F., Krammer, F., Baric, R.S., le Gars, M., Sadoff, J., de Groot, A.M., Heerwegh, D., Struyf, F., Douoguih, M., van Hoof, J., Schuitemaker, H., and Barouch, D.H. (2021) ‘Immunogenicity of Ad26.COV2.S vaccine against SARS-CoV-2 variants in humans’, Nature, 596(7871), 268–272, available: https://doi.org/10.1038/s41586-021-03681-2.

Baker, J.M., Nakayama, J.Y., O’Hegarty, M., McGowan, A., Teran, R.A., Bart, S.M., Mosack, K., Roberts, N., Campos, B., Paegle, A., McGee, J., Herrera, R., English, K., Barrios, C., Davis, A., Roloff, C., Sosa, L.E., Brockmeyer, J., Page, L., Bauer, A., Weiner, J.J., Khubbar, M., Bhattacharyya, S., Kirking, H.L., and Tate, J.E. (2022) ‘SARS-CoV-2 B.1.1.529 (Omicron) Variant Transmission Within Households - Four U.S. Jurisdictions, November 2021-February 2022.’, MMWR. Morbidity and mortality weekly report, 71(9), 341–346, available: https://doi.org/10.15585/mmwr.mm7109e1.

Bechtel, T.J. and Weerapana, E. (2017) ‘From structure to redox: The diverse functional roles of disulfides and implications in disease’, PROTEOMICS, 17(6), 1600391, available: https://doi.org/10.1002/pmic.201600391.

Cai, Y., Zhang, J., Xiao, T., Lavine, C.L., Rawson, S., Peng, H., Zhu, H., Anand, K., Tong, P., Gautam, A., Lu, S., Sterling, S.M., Walsh, R.M., Rits-Volloch, S., Lu, J., Wesemann, D.R., Yang, W., Seaman, M.S., and Chen, B. (2021) ‘Structural basis for enhanced infectivity and immune evasion of SARS-CoV-2 variants.’, Science (New York, N.Y.), 373(6555), 642–648, available: https://doi.org/10.1126/science.abi9745.

Cai, Y., Zhang, J., Xiao, T., Peng, H., Sterling, S.M., Walsh, R.M., Rawson, S., Rits-Volloch, S., and Chen, B. (2020) ‘Distinct conformational states of SARS-CoV-2 spike protein.’, Science (New York, N.Y.), 369(6511), 1586–1592, available: https://doi.org/10.1126/science.abd4251.

Cao, Y., Wang, J., Jian, F., Xiao, T., Song, W., Yisimayi, A., Huang, W., Li, Q., Wang, P., An, R., Wang, J., Wang, Y., Niu, X., Yang, S., Liang, H., Sun, H., Li, T., Yu, Y., Cui, Q., Liu, S., Yang, X., Du, S., Zhang, Z., Hao, X., Shao, F., Jin, R., Wang, X., Xiao, J., Wang, Y., and Xie, X.S. (2022) ‘Omicron escapes the majority of existing SARS-CoV-2 neutralizing antibodies’, Nature, 602(7898), 657–663, available: https://doi.org/10.1038/s41586-021-04385-3.

Carreño, J.M., Alshammary, H., Tcheou, J., Singh, G., Raskin, A.J., Kawabata, H., Sominsky, L.A., Clark, J.J., Adelsberg, D.C., Bielak, D.A., Gonzalez-Reiche, A.S., Dambrauskas, N., Vigdorovich, V., Alburquerque, B., Amoako, A.A., Banu, R., Beach, K.F., Bermúdez-González, M.C., Cai, G.Y., Ceglia, I., Cognigni, C., Farrugia, K., Gleason, C.R., van de Guchte, A., Kleiner, G., Khalil, Z., Lyttle, N., Mendez, W.A., Mulder, L.C.F., Oostenink, A., Rooker, A., Salimbangon, A.T., Saksena, M., Paniz-Mondolfi, A.E., Polanco, J., Srivastava, K., Sather, D.N., Sordillo, E.M., Bajic, G., van Bakel, H., Simon, V., and Krammer, F. (2022) ‘Activity of convalescent and vaccine serum against SARS-CoV-2 Omicron’, Nature, 602(7898), 682–688, available: https://doi.org/10.1038/s41586-022-04399-5.

Cele, S., Jackson, L., Khoury, D.S., Khan, K., Moyo-Gwete, T., Tegally, H., San, J.E., Cromer, D., Scheepers, C., Amoako, D.G., Karim, F., Bernstein, M., Lustig, G., Archary, D., Smith, M., Ganga, Y., Jule, Z., Reedoy, K., Hwa, S.H., Giandhari, J., Blackburn, J.M., Gosnell, B.I., Abdool Karim, S.S., Hanekom, W., Davies, M.A., Hsiao, M., Martin, D., Mlisana, K., Wibmer, C.K., Williamson, C., York, D., Harrichandparsad, R., Herbst, K., Jeena, P., Khoza, T., Kløverpris, H., Leslie, A., Madansein, R., Magula, N., Manickchund, N., Marakalala, M., Mazibuko, M., Moshabela, M., Mthabela, N., Naidoo, K., Ndhlovu, Z., Ndung’u, T., Ngcobo, N., Nyamande, K., Patel, V., Smit, T., Steyn, A., Wong, E., von Gottberg, A., Bhiman, J.N., Lessells, R.J., Moosa, M.Y.S., Davenport, M.P., de Oliveira, T., Moore, P.L., and Sigal, A. (2022) ‘Omicron extensively but incompletely escapes Pfizer BNT162b2 neutralization’, Nature, 602(7898), 654–656, available: https://doi.org/10.1038/s41586-021-04387-1.

Chen, R.E., Winkler, E.S., Case, J.B., Aziati, I.D., Bricker, T.L., Joshi, A., Darling, T.L., Ying, B., Errico, J.M., Shrihari, S., VanBlargan, L.A., Xie, X., Gilchuk, P., Zost, S.J., Droit, L., Liu, Z., Stumpf, S., Wang, D., Handley, S.A., Stine, W.B., Shi, P.Y., Davis-Gardner, M.E., Suthar, M.S., Knight, M.G., Andino, R., Chiu, C.Y., Ellebedy, A.H., Fremont, D.H., Whelan, S.P.J., Crowe, J.E., Purcell, L., Corti, D., Boon, A. C.M., and Diamond, M.S. (2021) ‘In vivo monoclonal antibody efficacy against SARS-CoV-2 variant strains’, Nature, 596(7870), 103–108, available: https://doi.org/10.1038/s41586-021-03720-y.

Cui, Z., Liu, P., Wang, N., Wang, L., Fan, K., Zhu, Q., Wang, K., Chen, R., Feng, R., Jia, Z., Yang, M., Xu, G., Zhu, B., Fu, W., Chu, T., Feng, L., Wang, Y., Pei, X., Yang, P., Xie, X.S., Cao, L., Cao, Y., and Wang, X. (2022) ‘Structural and functional characterizations of infectivity and immune evasion of SARS-CoV-2 Omicron.’, Cell, 185(5), 860–871.e13, available: https://doi.org/10.1016/j.cell.2022.01.019.

Dejnirattisai, W., Huo, J., Zhou, D., Zahradník, J., Supasa, P., Liu, C., Duyvesteyn, H.M.E., Ginn, H.M., Mentzer, A.J., Tuekprakhon, A., Nutalai, R., Wang, B., Dijokaite, A., Khan, S., Avinoam, O., Bahar, M., Skelly, D., Adele, S., Johnson, S.A., Amini, A., Ritter, T.G., Mason, C., Dold, C., Pan, D., Assadi, S., Bellass, A., Omo-Dare, N., Koeckerling, D., Flaxman, A., Jenkin, D., Aley, P.K., Voysey, M., Costa Clemens, S.A., Naveca, F.G., Nascimento, V., Nascimento, F., Fernandes da Costa, C., Resende, P.C., Pauvolid-Correa, A., Siqueira, M.M., Baillie, V., Serafin, N., Kwatra, G., da Silva, K., Madhi, S.A., Nunes, M.C., Malik, T., Openshaw, P.J.M., Baillie, J.K., Semple, M.G., Townsend, A.R., Huang, K.Y.A., Tan, T.K., Carroll, M.W., Klenerman, P., Barnes, E., Dunachie, S.J., Constantinides, B., Webster, H., Crook, D., Pollard, A.J., Lambe, T., Conlon, C., Deeks, A.S., Frater, J., Frending, L., Gardiner, S., Jämsén, A., Jeffery, K., Malone, T., Phillips, E., Rothwell, L., Stafford, L., Baillie, J.K., Openshaw, P.J., Carson, G., Alex, B., Andrikopoulos, P., Bach, B., Barclay, W.S., Bogaert, D., Chand, M., Chechi, K., Cooke, G.S., da Silva Filipe, A., de Silva, T., Docherty, A.B., dos Santos Correia, G., Dumas, M.E., Dunning, J., Fletcher, T., Green, C.A., Greenhalf, W., Griffin, J.L., Gupta, R.K., Harrison, E.M., Hiscox, J. A., Wai Ho, A.Y., Horby, P.W., Ijaz, S., Khoo, S., Law, A., Lewis, M.R., Liggi, S., Lim, W.S., Maslen, L., Merson, L., Meynert, A.M., Moore, S.C., Noursadeghi, M., Olanipekun, M., Osagie, A., Palmarini, M., Palmieri, C., Paxton, W.A., Pollakis, G., Price, N., Rambaut, A., Robertson, D.L., Russell, C.D., Sancho-Shimizu, V., Sands, C.J., Scott, J.T., Sigfrid, L., Solomon, T., Sriskandan, S., Stuart, D., Summers, C., Swann, O. v., Takats, Z., Takis, P., Tedder, R.S., Thompson, A.R., Thomson, E.C., Thwaites, R.S., Turtle, L.C., Zambon, M., Hardwick, H., Donohue, C., Griffiths, F., Oosthuyzen, W., Donegan, C., Spencer, R.G., Norman, L., Pius, R., Drake, T.M., Fairfield, C.J., Knight, S.R., Mclean, K. A., Murphy, D., Shaw, C.A., Dalton, J., Girvan, M., Saviciute, E., Roberts, S., Harrison, J., Marsh, L., Connor, M., Halpin, S., Jackson, C., Gamble, C., Plotkin, D., Lee, J., Leeming, G., Wham, M., Clohisey, S., Hendry, R., Scott-Brown, J., Shaw, V., McDonald, S.E., Keating, S., Ahmed, K.A., Armstrong, J.A., Ashworth, M., Asiimwe, I.G., Bakshi, S., Barlow, S.L., Booth, L., Brennan, B., Bullock, K., Catterall, B. W., Clark, J.J., Clarke, E.A., Cole, S., Cooper, L., Cox, H., Davis, C., Dincarslan, O., Dunn, C., Dyer, P., Elliott, A., Evans, A., Finch, L., Fisher, L.W., Foster, T., Garcia-Dorival, I., Gunning, P., Hartley, C., Jensen, R.L., Jones, C.B., Jones, T.R., Khandaker, S., King, K., Kiy, R.T., Koukorava, C., Lake, A., Lant, S., Latawiec, D., Lavelle-Langham, L., Lefteri, D., Lett, L., Livoti, L.A., Mancini, M., McDonald, S., McEvoy, L., McLauchlan, J., Metelmann, S., Miah, N.S., Middleton, J., Mitchell, J., Murphy, E.G., Penrice-Randal, R., Pilgrim, J., Prince, T., Reynolds, W., Ridley, P.M., Sales, D., Shaw, V.E., Shears, R.K., Small, B., Subramaniam, K.S., Szemiel, A., Taggart, A., Tanianis-Hughes, J., Thomas, J., Trochu, E., van Tonder, L., Wilcock, E., Zhang, J.E., Flaherty, L., Maziere, N., Cass, E., Carracedo, A.D., Carlucci, N., Holmes, A., Massey, H., Murphy, L., McCafferty, S., Clark, R., Fawkes, A., Morrice, K., Maclean, A., Wrobel, N., Donnelly, L., Coutts, A., Hafezi, K., MacGillivray, L., Gilchrist, T., Adeniji, K., Agranoff, D., Agwuh, K., Ail, D., Aldera, E.L., Alegria, A., Allen, S., Angus, B., Ashish, A., Atkinson, D., Bari, S., Barlow, G., Barnass, S., Barrett, N., Bassford, C., Basude, S., Baxter, D., Beadsworth, M., Bernatoniene, J., Berridge, J., Berry, C., Best, N., Bothma, P., Chadwick, D., Brittain-Long, R., Bulteel, N., Burden, T., Burtenshaw, A., Caruth, V., Chambler, D., Chee, N., Child, J., Chukkambotla, S., Clark, T., Collini, P., Cosgrove, C., Cupitt, J., Cutino-Moguel, M.T., Dark, P., Dawson, C., Dervisevic, S., Donnison, P., Douthwaite, S., Drummond, A., DuRand, I., Dushianthan, A., Dyer, T., Evans, C., Eziefula, C., Fegan, C., Finn, A., Fullerton, D., Garg, S., Garg, A., Gkrania-Klotsas, E., Godden, J., Goldsmith, A., Graham, C., Hardy, E., Hartshorn, S., Harvey, D., Havalda, P., Hawcutt, D.B., Hobrok, M., Hodgson, L., Hormis, A., Jacobs, M., Jain, S., Jennings, P., Kaliappan, A., Kasipandian, V., Kegg, S., Kelsey, M., Kendall, J., Kerrison, C., Kerslake, I., Koch, O., Koduri, G., Koshy, G., Laha, S., Laird, S., Larkin, S., Leiner, T., Lillie, P., Limb, J., Linnett, V., Little, J., Lyttle, M., MacMahon, M., MacNaughton, E., Mankregod, R., Masson, H., Matovu, E., McCullough, K., McEwen, R., Meda, M., Mills, G., Minton, J., Mirfenderesky, M., Mohandas, K., Mok, Q., Moon, J., Moore, E., Morgan, P., Morris, C., Mortimore, K., Moses, S., Mpenge, M., Mulla, R., Murphy, M., Nagel, M., Nagarajan, T., Nelson, M., Norris, L., O’Shea, M.K., Otahal, I., Ostermann, M., Pais, M., Panchatsharam, S., Papakonstantinou, D., Paraiso, H., Patel, B., Pattison, N., Pepperell, J., Peters, M., Phull, M., Pintus, S., Pooni, J.S., Planche, T., Post, F., Price, D., Prout, R., Rae, N., Reschreiter, H., Reynolds, T., Richardson, N., Roberts, M., Roberts, D., Rose, A., Rousseau, G., Ruge, B., Ryan, B., Saluja, T., Schmid, M.L., Shah, A., Shanmuga, P., Sharma, A., Shawcross, A., Sizer, J., Shankar-Hari, M., Smith, R., Snelson, C., Spittle, N., Staines, N., Stambach, T., Stewart, R., Subudhi, P., Szakmany, T., Tatham, K., Thomas, J., Thompson, C., Thompson, R., Tridente, A., Tupper-Carey, D., Twagira, M., Vallotton, N., Vancheeswaran, R., Vincent-Smith, L., Visuvanathan, S., Vuylsteke, A., Waddy, S., Wake, R., Walden, A., Welters, I., Whitehouse, T., Whittaker, P., Whittington, A., Papineni, P., Wijesinghe, M., Williams, M., Wilson, L., Winchester, S., Wiselka, M., Wolverson, A., Wootton, D.G., Workman, A., Yates, B., Young, P., Paterson, N.G., Williams, M.A., Hall, D.R., Fry, E.E., Mongkolsapaya, J., Ren, J., Schreiber, G., Stuart, D.I., and Screaton, G.R. (2022) ‘SARS-CoV-2 Omicron-B.1.1.529 leads to widespread escape from neutralizing antibody responses’, Cell, 185(3), 467–484.e15, available: https://doi.org/10.1016/j.cell.2021.12.046.

Dixon, A.S., Schwinn, M.K., Hall, M.P., Zimmerman, K., Otto, P., Lubben, T.H., Butler, B.L., Binkowski, B.F., Machleidt, T., Kirkland, T.A., Wood, M.G., Eggers, C.T., Encell, L.P., and Wood, K. v (2016) ‘NanoLuc Complementation Reporter Optimized for Accurate Measurement of Protein Interactions in Cells.’, ACS chemical biology, 11(2), 400–8, available: https://doi.org/10.1021/acschembio.5b00753.

Fenouillet, E., Barbouche, R., and Jones, I.M. (2007) ‘Cell entry by enveloped viruses: redox considerations for HIV and SARS-coronavirus.’, Antioxidants & redox signaling, 9(8), 1009–34, available: https://doi.org/10.1089/ars.2007.1639.

Fossum, C.J., Laatsch, B.F., Lowater, H.R., Narkiewicz-Jodko, A.W., Lonzarich, L., Hati, S., and Bhattacharyya, S. (2022) ‘Pre-Existing Oxidative Stress Creates a Docking-Ready Conformation of the SARS-CoV-2 Receptor-Binding Domain’, ACS Bio & Med Chem Au, 2(1), 84–93, available: https://doi.org/10.1021/acsbiomedchemau.1c00040.

Gobeil, S.M.-C., Henderson, R., Stalls, V., Janowska, K., Huang, X., May, A., Speakman, M., Beaudoin, E., Manne, K., Li, D., Parks, R., Barr, M., Deyton, M., Martin, M., Mansouri, K., Edwards, R.J., Eaton, A., Montefiori, D.C., Sempowski, G.D., Saunders, K.O., Wiehe, K., Williams, W., Korber, B., Haynes, B.F., and Acharya, P. (2022) ‘Structural diversity of the SARS-CoV-2 Omicron spike’, Molecular Cell, 82(11), 2050–2068.e6, available: https://doi.org/10.1016/j.molcel.2022.03.028.

Gobeil, S.M.-C., Janowska, K., McDowell, S., Mansouri, K., Parks, R., Stalls, V., Kopp, M.F., Manne, K., Li, D., Wiehe, K., Saunders, K.O., Edwards, R.J., Korber, B., Haynes, B.F., Henderson, R., and Acharya, P. (2021) ‘Effect of natural mutations of SARS-CoV-2 on spike structure, conformation, and antigenicity.’, Science (New York, N.Y.), 373(6555), available: https://doi.org/10.1126/science.abi6226.

Grishin, A.M., Dolgova, N. v., Landreth, S., Fisette, O., Pickering, I.J., George, G.N., Falzarano, D., and Cygler, M. (2022) ‘Disulfide Bonds Play a Critical Role in the Structure and Function of the Receptor-binding Domain of the SARS-CoV-2 Spike Antigen’, Journal of Molecular Biology, 434(2), 167357, available: https://doi.org/10.1016/j.jmb.2021.167357.

Iketani, S., Liu, L., Guo, Y., Liu, L., Chan, J.F.W., Huang, Y., Wang, M., Luo, Y., Yu, J., Chu, H., Chik, K.K.H., Yuen, T.T.T., Yin, M.T., Sobieszczyk, M.E., Huang, Y., Yuen, K.Y., Wang, H.H., Sheng, Z., and Ho, D.D. (2022) ‘Antibody evasion properties of SARS-CoV-2 Omicron sublineages’, Nature, 604(7906), 553–556, available: https://doi.org/10.1038/s41586-022-04594-4.

Jensen, K.S., Hansen, R.E., and Winther, J.R. (2009) ‘Kinetic and Thermodynamic Aspects of Cellular Thiol–Disulfide Redox Regulation’, Antioxidants & Redox Signaling, 11(5), 1047–1058, available: https://doi.org/10.1089/ars.2008.2297.

Khanna, K., Raymond, W.W., Jin, J., Charbit, A.R., Gitlin, I., Tang, M., Werts, A.D., Barrett, E.G., Cox, J.M., Birch, S.M., Martinelli, R., Sperber, H.S., Franz, S., Duff, T., Hoffmann, M., Healy, A.M., Oscarson, S., Pöhlmann, S., Pillai, S.K., Simmons, G., and Fahy, J. v. (2022) ‘Exploring antiviral and anti-inflammatory effects of thiol drugs in COVID-19’, American Journal of Physiology-Lung Cellular and Molecular Physiology, 323(3), L372–L389, available: https://doi.org/10.1152/ajplung.00136.2022.

Kim, S.J., Yao, Z., Marsh, M.C., Eckert, D.M., Kay, M.S., Lyakisheva, A., Pasic, M., Bansal, A., Birnboim, C., Jha, P., Galipeau, Y., Langlois, M.-A., Delgado, J.C., Elgort, M.G., Campbell, R.A., Middleton, E.A., Stagljar, I., and Owen, S.C. (2022) ‘Homogeneous surrogate virus neutralization assay to rapidly assess neutralization activity of anti-SARS-CoV-2 antibodies.’, Nature communications, 13(1), 3716, available: https://doi.org/10.1038/s41467-022-31300-9.

Lan, J., Ge, J., Yu, J., Shan, S., Zhou, H., Fan, S., Zhang, Q., Shi, X., Wang, Q., Zhang, L., and Wang, X. (2020) ‘Structure of the SARS-CoV-2 spike receptor-binding domain bound to the ACE2 receptor.’, Nature, 581(7807), 215–220, available: https://doi.org/10.1038/s41586-020-2180-5.

Lavillette, D., Barbouche, R., Yao, Y., Boson, B., Cosset, F.-L., Jones, I.M., and Fenouillet, E. (2006) ‘Significant redox insensitivity of the functions of the SARS-CoV spike glycoprotein: comparison with HIV envelope.’, The Journal of biological chemistry, 281(14), 9200–4, available: https://doi.org/10.1074/jbc.M512529200.

Lewnard, J.A., Hong, V.X., Patel, M.M., Kahn, R., Lipsitch, M., and Tartof, S.Y. (2022) ‘Clinical outcomes associated with SARS-CoV-2 Omicron (B.1.1.529) variant and BA.1/BA.1.1 or BA.2 subvariant infection in Southern California.’, Nature medicine, 28(9), 1933–1943, available: https://doi.org/10.1038/s41591-022-01887-z.

Liu, L., Iketani, S., Guo, Y., Chan, J.F.W., Wang, M., Liu, L., Luo, Y., Chu, H., Huang, Y., Nair, M.S., Yu, J., Chik, K.K.H., Yuen, T.T.T., Yoon, C., To, K.K.W., Chen, H., Yin, M.T., Sobieszczyk, M.E., Huang, Y., Wang, H.H., Sheng, Z., Yuen, K.Y., and Ho, D.D. (2022) ‘Striking antibody evasion manifested by the Omicron variant of SARS-CoV-2’, Nature, 602(7898), 676–681, available: https://doi.org/10.1038/s41586-021-04388-0.

Manček-Keber, M., Hafner-Bratkovič, I., Lainšček, D., Benčina, M., Govednik, T., Orehek, S., Plaper, T., Jazbec, V., Bergant, V., Grass, V., Pichlmair, A., and Jerala, R. (2021) ‘Disruption of disulfides within RBD of SARS-CoV-2 spike protein prevents fusion and represents a target for viral entry inhibition by registered drugs’, The FASEB Journal, 35(6), available: https://doi.org/10.1096/fj.202100560R.

Murae, M., Shimizu, Y., Yamamoto, Y., Kobayashi, A., Houri, M., Inoue, T., Irie, T., Gemba, R., Kondo, Y., Nakano, Y., Miyazaki, S., Yamada, D., Saitoh, A., Ishii, I., Onodera, T., Takahashi, Y., Wakita, T., Fukasawa, M., and Noguchi, K. (2022) ‘The function of SARS-CoV-2 spike protein is impaired by disulfide-bond disruption with mutation at cysteine-488 and by thiol-reactive N-acetyl-cysteine and glutathione.’, Biochemical and biophysical research communications, 597, 30–36, available: https://doi.org/10.1016/j.bbrc.2022.01.106.

Planas, D., Saunders, N., Maes, P., Guivel-Benhassine, F., Planchais, C., Buchrieser, J., Bolland, W.H., Porrot, F., Staropoli, I., Lemoine, F., Péré, H., Veyer, D., Puech, J., Rodary, J., Baele, G., Dellicour, S., Raymenants, J., Gorissen, S., Geenen, C., Vanmechelen, B., Wawina-Bokalanga, T., Martí-Carreras, J., Cuypers, L., Sève, A., Hocqueloux, L., Prazuck, T., Rey, F.A., Simon-Loriere, E., Bruel, T., Mouquet, H., André, E., and Schwartz, O. (2022) ‘Considerable escape of SARS-CoV-2 Omicron to antibody neutralization’, Nature, 602(7898), 671–675, available: https://doi.org/10.1038/s41586-021-04389-z.

Reynolds, C.J., Gibbons, J.M., Pade, C., Lin, K.-M., Sandoval, D.M., Pieper, F., Butler, D.K., Liu, S., Otter, A.D., Joy, G., Menacho, K., Fontana, M., Smit, A., Kele, B., Cutino-Moguel, T., Maini, M.K., Noursadeghi, M., Brooks, T., Semper, A., Manisty, C., Treibel, T.A., Moon, J.C., McKnight, Á., Altmann, D.M., and Boyton, R.J. (2022) ‘Heterologous infection and vaccination shapes immunity against SARS-CoV-2 variants’, Science, 375(6577), 183–192, available: https://doi.org/10.1126/science.abm0811.

Shang, J., Ye, G., Shi, K., Wan, Y., Luo, C., Aihara, H., Geng, Q., Auerbach, A., and Li, F. (2020) ‘Structural basis of receptor recognition by SARS-CoV-2.’, Nature, 581(7807), 221–224, available: https://doi.org/10.1038/s41586-020-2179-y.

Shi, Y., Zeida, A., Edwards, C.E., Mallory, M.L., Sastre, S., Machado, M.R., Pickles, R.J., Fu, L., Liu, K., Yang, J., Baric, R.S., Boucher, R.C., Radi, R., and Carroll, K.S. (2022) ‘Thiol-based chemical probes exhibit antiviral activity against SARS-CoV-2 via allosteric disulfide disruption in the spike glycoprotein.’, Proceedings of the National Academy of Sciences of the United States of America, 119(6), available: https://doi.org/10.1073/pnas.2120419119.

Suhail, S., Zajac, J., Fossum, C., Lowater, H., McCracken, C., Severson, N., Laatsch, B., Narkiewicz-Jodko, A., Johnson, B., Liebau, J., Bhattacharyya, S., and Hati, S. (2020) ‘Role of Oxidative Stress on SARS-CoV (SARS) and SARS-CoV-2 (COVID-19) Infection: A Review’, The Protein Journal, 39(6), 644–656, available: https://doi.org/10.1007/s10930-020-09935-8.

Viana, R., Moyo, S., Amoako, D.G., Tegally, H., Scheepers, C., Althaus, C.L., Anyaneji, U.J., Bester, P.A., Boni, M.F., Chand, M., Choga, W.T., Colquhoun, R., Davids, M., Deforche, K., Doolabh, D., du Plessis, L., Engelbrecht, S., Everatt, J., Giandhari, J., Giovanetti, M., Hardie, D., Hill, V., Hsiao, N.-Y., Iranzadeh, A., Ismail, A., Joseph, C., Joseph, R., Koopile, L., Kosakovsky Pond, S.L., Kraemer, M.U.G., Kuate-Lere, L., Laguda-Akingba, O., Lesetedi-Mafoko, O., Lessells, R.J., Lockman, S., Lucaci, A.G., Maharaj, A., Mahlangu, B., Maponga, T., Mahlakwane, K., Makatini, Z., Marais, G., Maruapula, D., Masupu, K., Matshaba, M., Mayaphi, S., Mbhele, N., Mbulawa, M.B., Mendes, A., Mlisana, K., Mnguni, A., Mohale, T., Moir, M., Moruisi, K., Mosepele, M., Motsatsi, G., Motswaledi, M.S., Mphoyakgosi, T., Msomi, N., Mwangi, P.N., Naidoo, Y., Ntuli, N., Nyaga, M., Olubayo, L., Pillay, S., Radibe, B., Ramphal, Y., Ramphal, U., San, J.E., Scott, L., Shapiro, R., Singh, L., Smith-Lawrence, P., Stevens, W., Strydom, A., Subramoney, K., Tebeila, N., Tshiabuila, D., Tsui, J., van Wyk, S., Weaver, S., Wibmer, C.K., Wilkinson, E., Wolter, N., Zarebski, A.E., Zuze, B., Goedhals, D., Preiser, W., Treurnicht, F., Venter, M., Williamson, C., Pybus, O.G., Bhiman, J., Glass, A., Martin, D.P., Rambaut, A., Gaseitsiwe, S., von Gottberg, A., and de Oliveira, T. (2022) ‘Rapid epidemic expansion of the SARS-CoV-2 Omicron variant in southern Africa’, Nature, 603(7902), 679–686, available: https://doi.org/10.1038/s41586-022-04411-y.

Walls, A.C., Park, Y.-J., Tortorici, M.A., Wall, A., McGuire, A.T., and Veesler, D. (2020) ‘Structure, Function, and Antigenicity of the SARS-CoV-2 Spike Glycoprotein.’, Cell, 181(2), 281–292.e6, available: https://doi.org/10.1016/j.cell.2020.02.058.

Wrapp, D., Wang, N., Corbett, K.S., Goldsmith, J.A., Hsieh, C.-L., Abiona, O., Graham, B.S., and McLellan, J.S. (2020) ‘Cryo-EM structure of the 2019-nCoV spike in the prefusion conformation.’, Science (New York, N.Y.), 367(6483), 1260–1263, available: https://doi.org/10.1126/science.abb2507.

Wu, C., Suzuki-Ogoh, C., and Ohmiya, Y. (2007) ‘Dual-reporter assay using two secreted luciferase genes.’, BioTechniques, 42(3), 290, 292, available: https://doi.org/10.2144/000112428.

Yao, Z., Drecun, L., Aboualizadeh, F., Kim, S.J., Li, Z., Wood, H., Valcourt, E.J., Manguiat, K., Plenderleith, S., Yip, L., Li, X., Zhong, Z., Yue, F.Y., Closas, T., Snider, J., Tomic, J., Drews, S.J., Drebot, M.A., McGeer, A., Ostrowski, M., Mubareka, S., Rini, J.M., Owen, S., and Stagljar, I. (2021) ‘A homogeneous split-luciferase assay for rapid and sensitive detection of anti-SARS CoV-2 antibodies.’, Nature communications, 12(1), 1806, available: https://doi.org/10.1038/s41467-021-22102-6.

Yin, W., Xu, Y., Xu, P., Cao, X., Wu, C., Gu, C., He, X., Wang, X., Huang, S., Yuan, Q., Wu, K., Hu, W., Huang, Z., Liu, J., Wang, Z., Jia, F., Xia, K., Liu, P., Wang, X., Song, B., Zheng, J., Jiang, H., Cheng, X., Jiang, Y., Deng, S.-J., and Xu, H.E. (2022) ‘Structures of the Omicron spike trimer with ACE2 and an anti-Omicron antibody.’, Science (New York, N.Y.), 375(6584), 1048–1053, available: https://doi.org/10.1126/science.abn8863.

Zhang, S., Go, E.P., Ding, H., Anang, S., Kappes, J.C., Desaire, H., and Sodroski, J.G. (2022) ‘Analysis of Glycosylation and Disulfide Bonding of Wild-Type SARS-CoV-2 Spike Glycoprotein.’, Journal of virology, 96(3), e0162621, available: https://doi.org/10.1128/JVI.01626-21.

